# AlphaPeptStats: an open-source Python package for automated, scalable and industrial-strength statistical analysis of mass spectrometry-based proteomics

**DOI:** 10.1101/2023.03.10.532057

**Authors:** Elena Krismer, Maximilian T Strauss, Matthias Mann

## Abstract

**Summary:** The widespread application of mass spectrometry (MS)-based proteomics in biomedical research increasingly requires robust, transparent and streamlined solutions to extract statistically reliable insights. Existing, popular tools were generally developed for specific uses in academic environments and did not fully embrace current open-source principles and best practices of software engineering. We have designed and implemented AlphaPeptStats, an inclusive python package with broad functionalities for normalization, imputation, visualization, and statistical analysis of proteomics data. It modularly builds on the established stack of Python scientific libraries, and is accompanied by a rigorous testing framework with 98% test coverage. It imports the output of a range of popular search engines. Data can be filtered and normalized according to user specifications. At its heart, AlphaPeptStats provides a wide range of robust statistical algorithms such as t-tests, ANOVA, PCA, hierarchical clustering and multiple covariate analysis – all in an automatable manner. Data visualization capabilities include heat maps, volcano plots, scatter plots in publication-ready format. AlphaPeptStats advances proteomic research through its robust tools that enable researchers to manually or automatically explore complex datasets to identify interesting patterns and outliers.

**Availability:** AlphaPeptStats is implemented in Python and part of the AlphaPept framework. It is released under a permissive Apache license. The source code and one-click installers are freely available and on GitHub at https://github.com/MannLabs/alphapeptstats.

## 1 Introduction

Mass spectrometry (MS)-based proteomics has emerged as a powerful tool in biomedical research (Aebersold and Mann, 2016). The rapid development of platforms and algorithms allows the identification and quantification of proteins with ever greater depth and precision. These workflows and search engines produce tables of identified and quantified proteins, which then require rigorous statistical methods to identify robust patterns and potentially biologically interesting outliers.

To date, we and others have developed popular applications, such as MSstats (Choi et al., 2014), Perseus (Tyanova *et al*., 2016) and MSPypeline (Heming et al., 2022) for the downstream analysis of proteomics data. While these tools mostly cover the required steps in the analysis pipeline, they can be limited in the search engines they support, access to the source code, test coverage, automation, and the ability to easily implement the latest algorithms. Furthermore, some of their functionality can readily be leveraged by domain experts, but this is more challenging for non-experts who need to integrate biological knowledge and contextualize the findings. This constitutes the need for an easy-to-use, rigorous and robust tool to maximize the biological insight that can be extracted from proteomics data.

## 2 The AlphaPeptStats library

As part of our recently developed AlphaPept framework (Strauss *et al*., 2021; Voytik *et al*., 2022; Zeng *et al*., 2022), we implemented AlphaPeptStats in Python because of its straightforward syntax and the availability of high-quality scientific libraries. AlphaPeptStats is built on top of highly performant, widely used and community-tested computing packages such as NumPy (Harris *et al*., 2020), Plotly (Plotly Technologies Inc., 2015), Pandas (McKinney, 2010) and SciPy (Virtanen *et al*., 2020). We additionally implemented state-of-the-art bioinformatic libraries, such as *diffxpy* from the *Scanpy-package* for differential expression analysis (Wolf *et al*., 2018) and *a GO tool* for enrichment analysis with gene ontology (GO)-terms, tailored for MS (Schölz *et al*., 2015).

The AlphaPeptStats source code is freely available on GitHub under the permissive Apache license. The package can readily be installed from PyPI using the standard pip module. Additionally, we provide one-click installers for Linux, MacOS and Windows and a Dockerfile for containerized deployment, e.g., in cloud environments. Furthermore, automated postprocessing workflows can be created in AlphaPeptStats. This can also be used to systematically iterate over available options such as different normalization methods to identify best-performing ones.

Additionally, we have deployed a web-based version of AlphaPeptStats that is hosted on Streamlit-share (see GitHub repository for the link) (Snowflake Inc., 2023). This enables users to explore and use AlphaPeptStats without requiring the installation of any software.

We designed AlphaPeptStats with best software engineering practices in mind, including continuous integration pipelines on GitHub, ensuring that the software is continuously tested and verified. Our extensive testing framework reports a test coverage of 98%, providing confidence in accuracy and reliability of the software, in line with standard packages such as NumpPy or Pandas.

Extensive documentation of the AlphaPeptStats functionalities was a key part of this project and include several Jupyter notebooks that serve as tutorials to guide novice users. These notebooks are designed to encourage user engagement and offer a step-by-step approach to learning the package, Alternatively, the graphical user interface is straightforward to learn as well as allowing quick and easy data exploration.

## 3 Overview of the AlphaPeptStats workflow

At present, AlphaPeptStats is already capable of importing and processing proteomics data generated from multiple software platforms, including MaxQuant (Cox and Mann, 2008), AlphaPept, DIA-NN (Demichev *et al*., 2020), Spectronaut (Bruderer *et al*., 2015), and the FragPipe computational framework (Kong *et al*., 2017; Teo *et al*., 2021; Yu *et al*., 2020, 2021; da Veiga Leprevost *et al*., 2020) (Fig. 1A). The modular design of the import functions allow straightforward extensions to other data formats.

**Fig 1:**
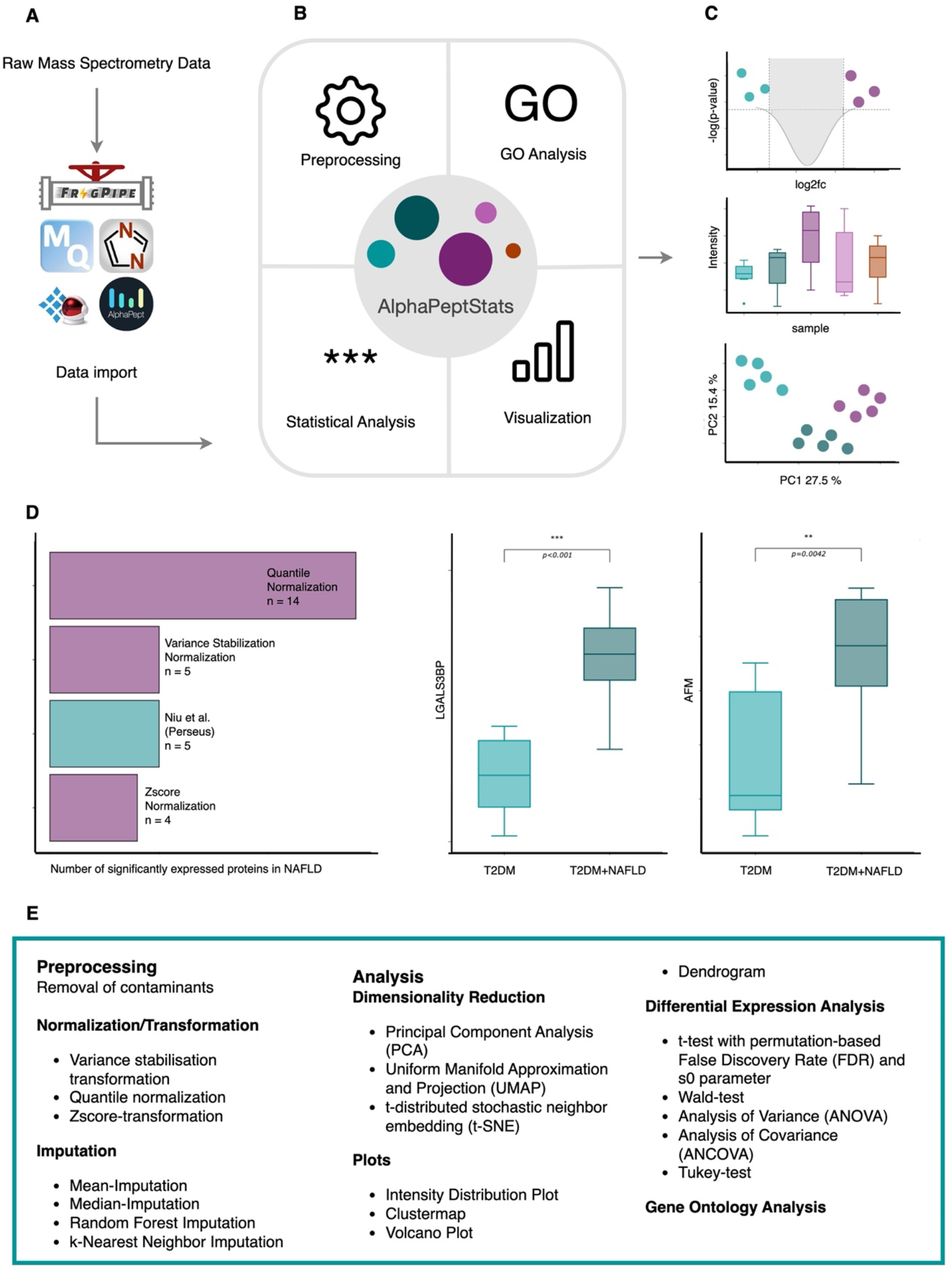
A AlphaPeptStats workflow. Output files from FragPipe, MaxQuant, DIA-NN, Spectronaut and AlphaPept can be imported. B The package includes functionalities for preprocessing, statistical analysis, visualization, and gene ontology (GO) term analysis C Figures can be exported as vector graphics for subsequent use in publications. D Systematic optimization of several normalization methods on a NAFLD dataset leading to different numbers of significantly expressed proteins with up to 14 when using Quantile Normalization. E Example plots of LGALS3BP and AFM showing significant expression F Table of processing steps that are available in AlphaPeptStats. Abbreviations: T2DM: Diabetes-Mellitus Type 2, NAFLD: non alcoholic fatty liver disease

Users are required to specify their proteomics results file and accompanying metadata. AlphaPeptStats provides a high-level API by storing data in a Python class named DataSet, with multiple methods ranging from data preprocessing, statistical analysis, GO analysis, to visualization (Fig. 1B.). The latter can export vector graphics for subsequent use in publications (Fig. 1C). An overview of the processing steps in AlphaPeptStats is provided in Figure 1D.

### 3.1 Preprocessing

After loading the data into a DataSet object, the user can select multiple optional preprocessing steps ranging from the removal of contaminants, normalization to imputation. For contaminant removal, AlphaPeptStats uses a recently developed library (Frankenfield *et al*., 2022) to help decrease false discoveries. In addition, AlphaPeptStats incorporates various normalization and imputation techniques to facilitate robust and accurate data analysis. One of the methods that we integrated - random forest imputation - has demonstrated superior performance compared to other commonly used imputation methods in several studies (Jin *et al*., 2021; Kokla *et al*., 2019). This algorithm was imported from scikit-learn, demonstrating how easily state of the art methods can be added to AlphaPeptStats.

Importantly, all selected preprocessing steps are stored in the DataSet object, ensuring reproducibility. Different normalization and imputation methods can be systematically assessed as AlphaPeptStats can iterate through them automatically by means of passing a single parameter to the plotting functions.

### 3.2 Visualization and Statistical Analysis

Users can visualize their results via dedicated functions that allow the straightforward interpretation of the data, including principal component analysis (PCA) plots, heatmaps, dendrogram and volcano plots. Figures can be exported as publication-ready scalable vector graphics. AlphaPeptStats leverages the capabilities of the Plotly graphing library, producing interactive and zoomable graphs by default and enabling advanced users to tailor the generated figures to their specific needs and preferences.

Statistical testing for differential expression analysis can be performed using Analysis of Variance (ANOVA), Analysis of Covariance (ANCOVA) or t-testing. We further provide a reimplementation from R of the significance analysis of microarrays (SAM), which is a very widely used algorithm in proteomic (Tusher *et al*., 2001). Significantly expressed proteins can then be subjected to GO annotation.

### 3.3 Graphical User Interface

As AlphaPeptStats is a Python library it can be imported and used in any Python program, scripts or Jupyter Notebooks. As mentioned, figures are produced by the incorporated Plotly library, making graphs interactively explorable.

Furthermore, the popular Streamlit library provides even easier access to AlphaPeptStats functionalities and output for non-coders. In this case, the graphical user interface of AlphaPeptStats enables users to directly select functions and analyze their data in a browser-based environment.

## 4 Application of AlphaPeptStats

To exemplify the capabilities of AlphaPeptStats, we applied it to our recently published study on non-alcoholic liver disease (NAFLD) (Niu *et al*., 2019). Using default settings, this successfully reproduced the same significantly differentially expressed proteins as previously reported, except for one (PIGR). As PIGR was validated as a biomarker in that study, we investigated the reason for its disappearance. We found that this biomarker was just below our stringent cutoff criteria due to too many missing values. Next, we leveraged the automatic parameter optimization and systematically investigated how the different preprocessing steps would maximize the number of differentially expressed proteins. This uncovered 9 additional biomarkers for NAFLD compared to the original five when applying quantile normalization, shown in Fig. 1E. Exploration of the additional biomarkers indicated a plausible connection to NAFLD (Suppl. Table 1). Exemplary plots generated with AlphaPeptStats of the enriched proteins LGALS3BP and AFM in NAFLD are presented in Figure 1F.

## 5 Conclusion

We developed AlphaPeptStats, a user-friendly, open-source package dedicated to the downstream analysis of mass spectrometry data, covering all steps from preprocessing and statistical analysis to visualization. Apart from stand-alone use, it can also easily be incorporated into automated bioinformatics pipelines. It features extensive tests and robust design principles of software engineering on GitHub, such as continuous testing and continuous integration to ensure a stable and reliable workflow. Its modular framework allows extensions with additional functionality. Leveraging its ability to programmatically test different normalization methods we more than doubled differentially expressed potential biomarkers in NAFLD compared to our previously published analysis. We envision that AlphaPeptStats will be a suitable standard for statistical analysis and exploration for the challenging proteomics data set that can readily be produced today.

## Author contributions

M.T.S., E.K. and M.M. conceptualized the project. E.K. implemented and evaluated the AlphaPeptStats functions with the support of M.T.S.. All authors wrote the manuscript.

## Acknowledgements

The authors thank Frederik Post, Sonja Kabatnik, Andreas Mund, Lili Niu and our colleagues at the MPI and CPR for testing and providing critical feedback on AlphaPeptStats.

