## Supplementary Material for "AlphaPeptStats: an open-source Python package for automated, scalable and industrial-strength statistical analysis of mass spectrometry-based proteomics"

| Protein | Full name | Biological Process | Supporting Literature |
|---|---|---|---|
| C9 | Complement component C9 | Immune system regulation and inflammation | (Subudhi et al., 2022) |
| SERPINC1 | Serpin peptidase inhibitor, clade C (antithrombin), member 1 | Coagulation cascade | (Bell et al., 2010) |
| ANPEP | Alanyl aminopeptidase | Angiogenesis | (Martinou et al., 2022) |
| ITIH4 | Inter-alpha-trypsin inhibitor heavy chain 4 | Immune system regulation and inflammation | (Martinou et al., 2022) |
| APOB | Apolipoprotein B | Cholesterol and triglyceride balance | (Bell et al., 2010) |
| C3 | Complement component C3 | Immune system regulation and inflammation | (Ogresta et al., 2022) |
| FGA | Fibrinogen A alpha | Coagulation cascade | (Ogresta et al., 2022) |
| FGG | Fibrinogen gamma chain | Coagulation cascade | (Ogresta et al., 2022) |
| AMBP | Alpha-1-Microglobulin/Bikunon | Host-virus interaction | (Berezin et al., 2023) |
| CFH | Complement factor H | Immune system regulation and inflammation | (Ogresta et al., 2022) |
| SERPINF2 | Serpin Family F member 2 | Coagulation cascade | (Bell et al., 2010) |
| AMF | Autocrine motility factor | Gluconeogenesis, Glycolisis | (Lin et al., 2020) |
| 1TP5BP1 | Tumor Protein P53 Binding Protein 1 | DNA damage, repair | (Akazawa et al., 2019) |
| GPLD1 | Glycosylposphatidylinositol sepcific phospholipase D1 | Lipid metabolism | (Yuan et al., 2008) |
| LGALS3BP | lectin | Cell adhesion | (Wood et al., 2017) |

Supplementary Table 1. Significantly altered proteins in non-alcoholic fatty liver disease (NAFLD) , identified with AlphaPeptStats.
